# Identification of a novel interaction between *Theileria* Prohibitin (*Ta*PHB-1) and bovine RUVBL-1

**DOI:** 10.1101/2022.02.14.480320

**Authors:** Prasanna Babu Araveti, Prajna Parimita Kar, Akshay Kuriakose, Achintya Sanju, Anand Srivastava

**Author notes:** Corresponding author: Anand Srivastava, National Institute of Animal Biotechnology (NIAB), Gachibowli, Gopanpalli, Hyderabad, Telangana 500 032, India.

## Abstract

Bovine tropical theileriosis causes huge economic loss worldwide. It is a tick borne disease of bovine caused by the parasite *Theileria annulata. T. annulata* is an intracellular parasite that belongs to the phylum Apicomplexa. The sporozoites of *T. annulata* are released by the tick into the bloodstream of the host during the blood meal that invades bovine B cells, macrophages, or monocytes. This infection leads to the transformation of the host cells and brings cancer-like phenotype in the host cells. The parasite proteins play a vital role in the transformation of the host cell. However, the parasite factors involved in the host cell transformation are not well explored. Previously, *Ta*PIN1, a peptidyl-prolyl isomerase of *T. annulata*, was shown to be secreted to the host cytosol and play a role in the host cell transformation. The present study was carried out to explore the parasite-host interactions that may play an important role in the host cell transformation. We identified the parasite proteins that are expressed in the schizont stage with a signal peptide. We narrow down our search to a parasite prohibitin. The *in silico* analysis of *T. annulata* prohibitin (TA04375, *Ta*PHB-1) showed that *Ta*PHB-1 shares homology with the mammalian prohibitin 1. With the localization experiments, we confirmed that *Ta*PHB-1 is exported to the parasite surface and also to the host cell cytosol. Further, we observed that the localization of host prohibitin differs in the parasite-infected cells and could not be reverted back by the elimination of the parasite in the infected cells. We found through the yeast-two-hybrid studies that bovine RUVBL1 (BoRUVBL-1) interacts with *Ta*PHB-1. The interaction between BoRUVBL1 and *Ta*PHB-1 was predominantly observed on the parasite surface in the infected bovine cells. The interaction was further confirmed with immunoprecipitation and LC-MS/MS analysis. Further, the LC-MS/MS based *Ta*PHB-1 interactome study reveals that it interacts with proteins that regulate actin cytoskeleton organization, protein folding, mRNA processing, and metabolic processes. Our finding suggests that the parasite releases prohibitin protein into the cytoplasm of the host cell where it interacts with the host RUVBL-1. This finding has implications not only in the understanding of *Theileria* parasite biology in greater depth but also in the cancer biology where previously differential localization of prohibitin proteins was observed but its interaction partner was not known.

**Author summary:** *Theileria annulata*, an apicomplexan, is a unique parasite which can transform host leucocytes. This parasite uses this strategy for its own multiplication. The cells infected with this parasite, when treated with buparvaquone, an anti-theilerial drug, cannot survive without the parasite. This observation suggests that the parasite derived factors are required to maintain the cancerous phenotype of the host cell. We mined the parasite proteome to find out the proteins with signal sequence that may be secreted to the host cell cytosol and being expressed in the schizont stage. The parasite prohibitin (*Ta*PHB-1) chosen for this study was found to be secreted to the host cytoplasm and on the parasite surface. Interestingly, we observed a noticeable change in the localization of the host prohibitin in the parasite infected cells. The host prohibitin that is normally localized to the mitochondria in the uninfected cells was observed in the host cell nucleus similar to the cancerous cells. Since the parasite protein is exported to the host cytoplasm we looked for its interacting partner. We performed yeast-two-hybrid screening with *Ta*PHB-1 with in-house prepared the cDNA library of the infected bovine leucocytes. We identified bovine RUVBL1 as the interacting partner of *Ta*PHB-1. Interestingly, the interaction between parasite prohibitin and bovine RUVBL1 was observed on the parasite surface. Further, analysis of the parasite prohibitin interactome in the infected cells shows that it might be involved with those proteins which regulate actin cytoskeleton organization, protein folding, mRNA processing and metabolic process. Since parasite infected cells have cancer like phenotype, the identification of this novel interaction will open up new avenues not only in the arena of parasite biology but also in the domain of cancer biology.

## Introduction

Tropical theileriosis or Mediterranean fever is a tick-borne haemoprotozoan disease caused by *Theileria annulata*, an intracellular parasite. Tropical theileriosis is geographically distributed in various countries in Central Asia, Southern Europe, North Africa and Middle East. *T. annulata* is transmitted by *Hyalomma anatolicum*. It causes disease in *Bos taurus* and *Bubalus bubalis* (1). The symptoms of the disease include fever, enlarged peripheral lymph nodes, anemia, decline in milk production and emaciation. Death may occur after 3-4 weeks of infection in untreated animals. The economic impact of tropical theileriosis in India is approximately US$ 800 million (2). Buparvaquone, a hydroxynaphthoquinone, is effective in the treatment of bovine tropical theileriosis (3). Point mutations in *T. annulata* cytochrome b (4) and alanine-to-proline mutation at position 53 of *Ta*PIN1 (5) leads to the gain of resistance towards buparvaquone in this parasite. The complex life cycle of *T. annulata* includes gametogony and sporogony in tick while merogony and piroplasm stage in cattle. The cattle is infected by the inoculation of sporozoites by the tick during the feeding of blood meal. The sporozoites invade leucocytes (B-cells and monocytes) and develop into the macroschizonts. The macroschizonts develop into microschizonts and ultimately into the merozoites which evade from the leucocytes. The merozoite infects erythrocytes where it develops into piroplasm. The piroplasms are taken up by the tick and they undergo gametogony in gut epithelial cells of the tick and finally develop into sporozoites by sporogony in salivary glands.

The macroschizont is one of the most important stages as macroschizonts can subvert various cell signaling pathways of the host cell which lead to the transformation of the host leucocytes (6). The parasite proteins play a vital role in subverting the cell signaling pathways as killing of the parasite leads to the death of the host cells (7). The interaction between host and parasite protein may be involved in the transformation of the host cell. The parasite peptidyl-prolyl isomerase (*Ta*PIN1) is secreted into host cytosol and interacts with host ubiquitin ligase FBW7, leading to its degradation and subsequent stabilization of c-JUN, promotes transformation (8). *T. annulata* p104 protein (*Ta*-p104) along with a microtubule and SH3 domain-interacting protein (*Ta*MISHIP) interacts with the host adaptor protein CD2AP, which may play a crucial role in cytokinesis as the overexpression of *Ta*MISHIP in non-infected bovine macrophages leads to binucleation (9). Also, the secreted *T. annulata Ta*9 protein contributes to the activation of the host AP-1 transcription factor and contributes to leucocyte transformation (10). Some other host-parasite interactions were reported which may not be involved in the host cell transformation but may be important for the parasite survival within the host. *T. annulata* cysteine proteinase interacts with two host proteins, cereblon transcript variant X2, and protein phosphatase 4 catalytic subunit which are involved in cellular processes like microtubule organization, DNA repair and cell apoptosis (11). *T. annulata* cyclophilin1 interacts with host cell MED21 which is normally involved in regulating the transcription of RNA polymerase II-dependent genes but knockdown of MED21 in *T. annulata*-infected leucocytes had no effect on NF-κB signaling (12). *T. annulata* surface protein (*Ta*SP) interacts with host microtubulin (13). During host cell mitosis *Ta*SP co-localizes and interacts with the spindle poles, which suggests that this interaction has a potential role in parasite distribution into the host cells (13). Also, *Ta*SP is phosphorylated by the host cell kinase CDK1 which plays a crucial role in cell division (14).

Prohibitins are highly conserved proteins found in all eukaryotes. They belong to stomatin/prohibitin/flotillin/HfIK/C (SPFH) family. They play a significant role in transcription, nuclear signaling, mitochondrial structural integrity, cell division and cell membrane metabolism (15). Distinct functions of prohibitin protein have been observed in various parasites. For example, prohibitin in *Trypanosoma brucei* is located in the mitochondrion where it is involved in the maintenance of mitochondrial membrane potential (16). Prohibitin in *Leishmania donovani* interacts with the host HSP70 present on macrophage surface which may help the parasite to escape invasion by host macrophages (17). In the case of *Plasmodium berghei*, prohibitin regulates mitochondrial membrane polarity and prohibitin deficient parasite causes mitochondrial depolarization in the mosquito vector, which, in turn, leads to a block in transmission (18). The role of prohibitin in the *T. annulata* parasite is unknown. However, previously it was speculated that prohibitin may have a role in transformation of the host cell (19). In this study, we characterized the prohibitin of *T. annulata* and identified its interacting partner.

## Results

### Prohibitins are highly conserved proteins

The screening of *T. annulata* proteome for proteins possessing a signal peptide using SignalP3.0 led to the identification of 438 proteins (S1 sheet). Forty nine out of these 438 proteins were found to be expressed in the schizont stage based on the available experimental *T. annulata* schizont proteome data (20, 21). Only 25 of these 49 proteins were non-hypothetical (S1 Fig and S1 sheet). Our analysis is consistent with the previously published data (19). We selected prohibitin(s) out of these 25 proteins for further analysis. The phylogenetic analysis of prohibitin proteins suggests that the prohibitins of phylum Apicomplexa branch away from the higher eukaryotes such as *Bos taurus* and *Homo sapiens* (Fig 1A). Among apicomplexans, the *T. annulata* prohibitins are closely related to *Babesia bovis* and distantly related to *Cryptosporidium parvum*. We also observed the presence of an additional prohibitin (PHB-3) in apicomplexan parasites except in *Cryptosporidium parvum*. Based on the phylogenetic tree analysis the three putative prohibitins of *T. annulata* were named as; TA04375 (*Ta*PHB-1), TA19320 (*Ta*PHB-2) and TA08975 ((*Ta*PHB-3) prohibitin like protein). The multiple sequence alignment and the identity matrix of these three prohibitins of *T. annulata* suggest that *Ta*PHB-3 is the most divergent prohibitin (Fig S2A and S2B). The subcellular localization prediction by the CELLO server of three putative prohibitin proteins of *T. annulata* suggests that *Ta*PHB-1 possesses a putative signal sequence that could localize this protein to the cytoplasm of the host while other two prohibitins were predicted to be localized in the mitochondria/chloroplast (Fig 1B). Thus, we hypothesized that if *Ta*PHB-1 protein is transported to the cytoplasm of the host then it could interact with the host protein(s) and may help in the transformation of the host cell. Hence, we selected *Ta*PHB-1 for further analysis.

**Fig 1.**
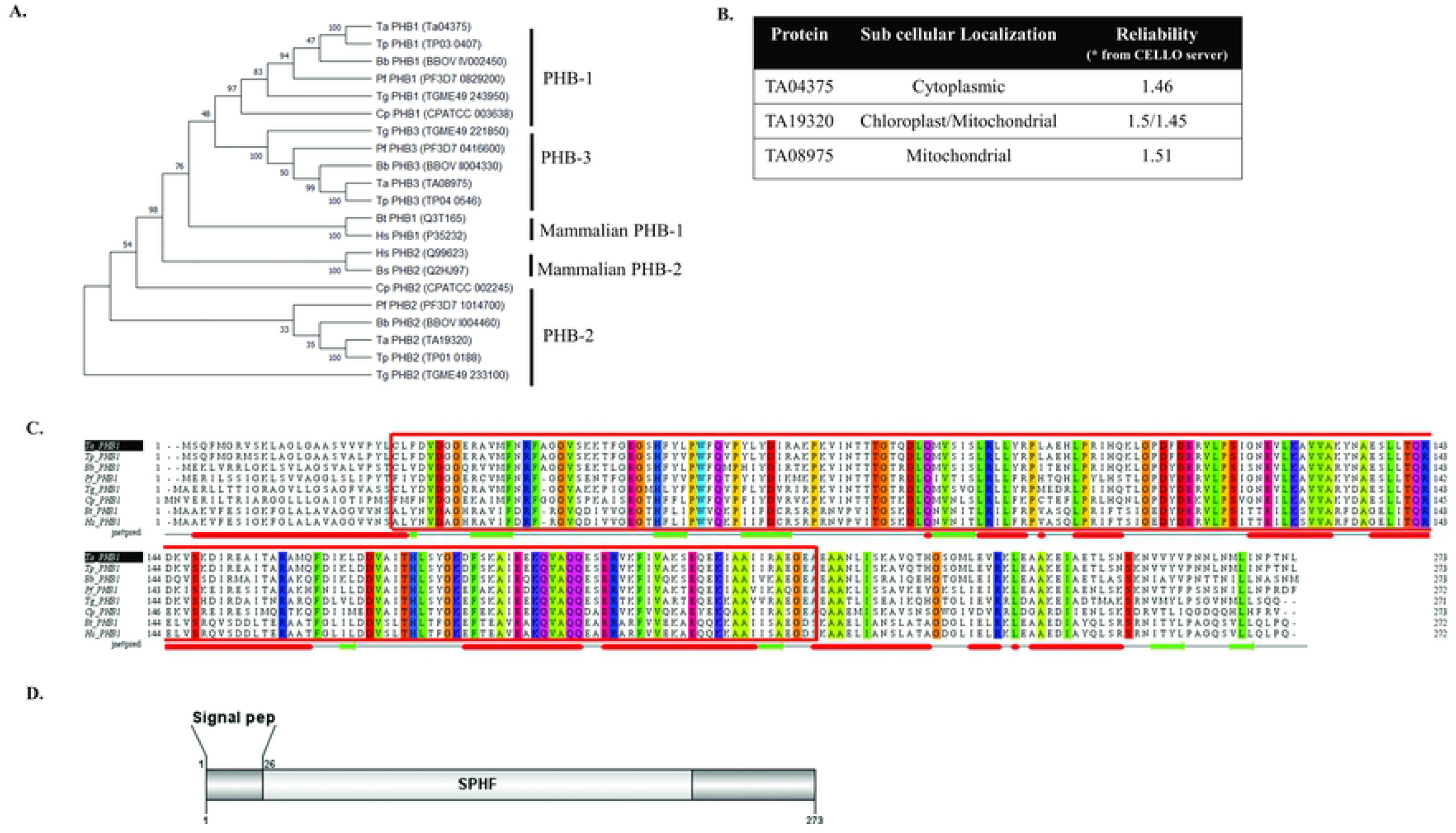
*In-silico* analysis of Prohibitins of *T. annulata*. **A.** Phylogenetic tree showing that *T.annulata* prohibitins (*Ta*PHB1, *Ta*PHB2 and *Ta*PHB3) are more close to *Theileria parva, Babesia bovis*. *Ta*PHB1 and *Ta*PHB2 are distantly related to *Cryptosporidium parvum*. PHB3 is absent in *Cryptosporidium parvum* and in higher eukaryotes like *Bos taurus* and *Homo sapiens*. Also, it is observed that *Ta*PHB1 and *Ta*PHB2 are distantly related to PHB2 of *Bos taurus* and *Homo sapiens*. **B.** Subcellular localization prediction of *T. annulata* prohibitins using CELLO shows that TA04375 may localize in the cytoplasm whereas others in the mitochondria. **C.** Multiple sequence alignment of *Ta*PHB1 (TA04375) with PHB1 of other apicomplexan parasites, *Bos taurus* and *Homo sapiens*, highlighted the SPHF domain with a red color box and the amino acid having 100% identity are shown in color. **D.** Domain analysis of *Ta*PHB1 containing SPHF domain and signal peptide.

Further, the multiple sequence alignment of protein sequences of Prohibitin-1 (PHB-1) of apicomplexan parasites (*Theileria annulata, Theileria parva, Plasmodium falciparum, Toxoplasma gondii, Cryptosporidium parvum, Babesia bovis*), human (*Homo sapiens*) and bovine (*Bos taurus*) suggested that these proteins are highly conserved and possess SPHF domain with a consensus sequence KQVAQQ (Fig 1C). The domain analysis of *Ta*PHB-1 is shown in fig 1D.

### Affinity purified *Ta*PHB-1 antibodies are specific to Ana2014 cells

*T. annulata* surface protein (*Ta*SP) is widely used as a cell surface marker for *T. annulata* schizonts (22). Thus, we expressed and purified recombinant *Ta*SP. The *Ta*SP protein is known to have an abnormal separation on SDS-PAGE (22). The expected molecular weight of cloned and expressed recombinant *Ta*SP is 15.4 kDa while we observed that it migrates at 36 kDa on SDS-PAGE (Fig S3). The antibodies raised in chicken against *Ta*SP could recognize native protein in the parasite lysate and also were able to stain parasite cell surface (S3 Fig and Fig 2C).

**Fig 2.**
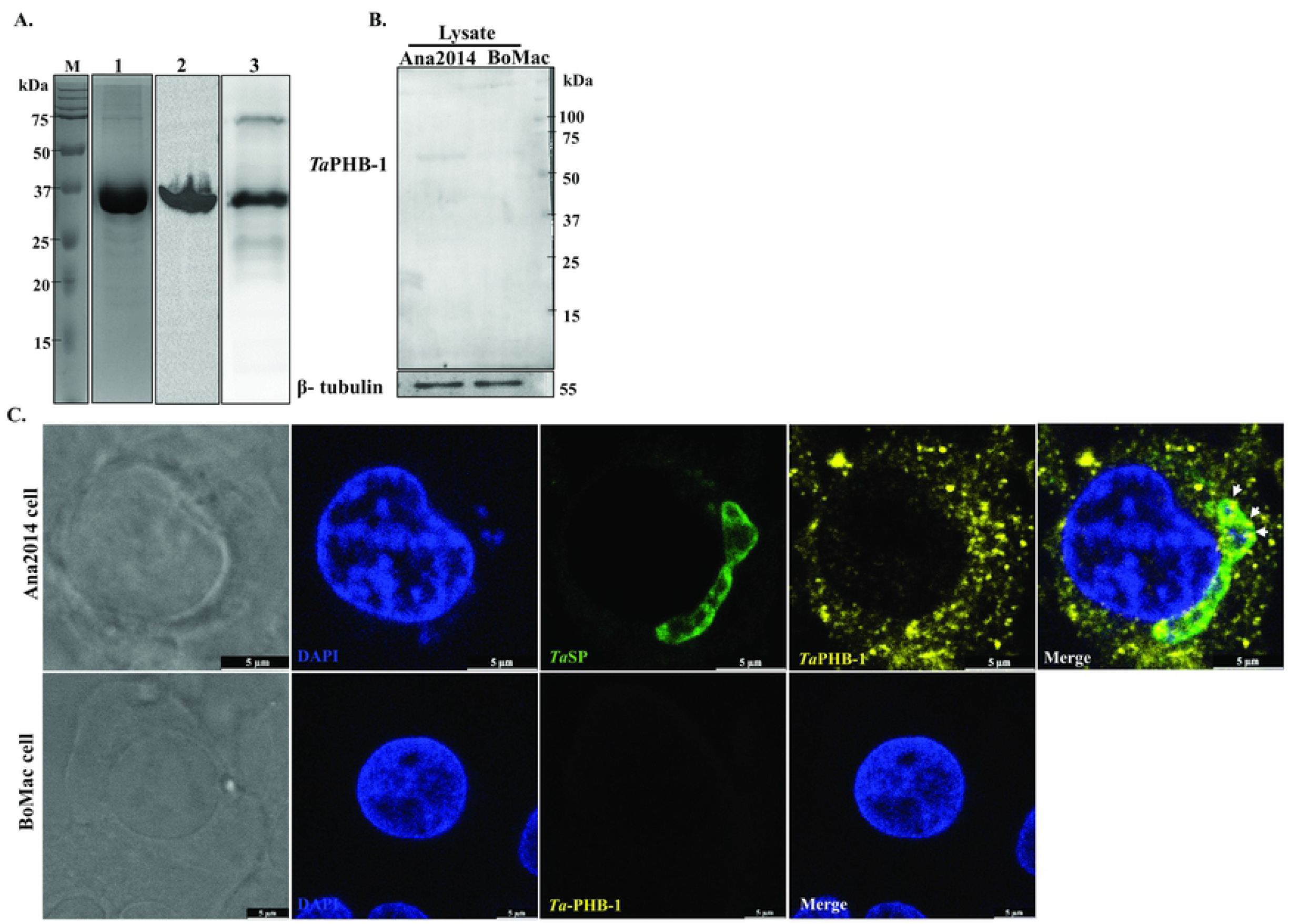
Affinity purification of anti-*Ta*PHB-1 mice sera and subcellular localization of *Ta*PHB-1 in Ana2014 cells. **A.** Protein expression, purification, and generation of mice polyclonal antibodies of *Ta*PHB-1. His tagged *Ta*PHB-1 protein purification from inclusion bodies under denatured conditions using Ni-NTA agarose beads. M: Marker. 1: Coomassie brilliant blue stained gel-elute with pH-4.3 buffer, 2: Western blotting of purified *Ta*PHB-1 probed with His-tag antibody, 3: Western blotting of purified *Ta*PHB-1 probed with anti-*Ta*PHB-1 mice sera. **B.** Western blot analysis of affinity purified *Ta*PHB-1 polyclonal antibodies against Ana2014 and BoMac cell lysates, β–tubulin was used as loading control. **C**. Ana2014 and BoMac cell lysates immuno-stained with affinity purified *Ta*PHB-1, *Ta*SP antibodies, BoMac cells were used as control. Scale bar-5 μm.

All our efforts to express *Ta*PHB-1 as a soluble protein remained unsuccessful. Thus, we purified recombinant His-tagged *Ta*PHB-1 under denatured condition (Fig 2A). The mice sera raised against denatured *Ta*PHB-1 detect recombinant His-tagged *Ta*PHB-1 in western blot analysis (Fig 2A). The mice sera against *Ta*PHB-1 were affinity purified against recombinant His-tagged *Ta*PHB-1. The expected size of *Ta*PHB-1 is 30kDa. In the western blot analysis, the affinity purified antibodies against *Ta*PHB-1 could recognize *Ta*PHB-1 protein at a higher molecular size (~60kDa) in the lysates of Ana2014 cells (Fig 2B). The LC-MS/MS analysis of recombinant *Ta*PHB-1 confirmed that the protein used for raising antibodies was *Ta*PHB-1 only (S4 Fig). Further, the affinity purified antibodies against *Ta*PHB-1 could not recognize any protein in the BoMac cell lysate (Fig 2B). As the signal was observed only from Ana2014 cell lysate in western blotting, we concluded that the affinity purified antibodies were specific to *Ta*PHB1 and did not cross react with BoPHB-1.

### *Ta*PHB-1 is exported out to the host cytoplasm by the parasite

As mentioned earlier, the signal sequence analysis of the prohibitins of *T. annulata* suggests that *Ta*PHB-1 has a signal sequence that may export this protein to the host cytoplasm. We performed confocal microscopy with affinity purified antibodies against *Ta*PHB-1 to localize this protein in the infected cells. The parasite surface was labelled with *Ta*SP antibodies. The *Ta*PHB-1 protein was found to be localized in the host cell cytosol (Fig 2C). We also observed *Ta*PHB-1 protein on the parasite surface (Fig 2C). We could not observe any signal from the BoMac cells when probed with affinity purified *Ta*PHB-1 antibodies. This suggests that the *Ta*PHB-1 protein can cross the parasite cell membrane to reach to the host cytoplasm.

### Localization of Bovine prohibitin-1 changes in the *T. annulata* infected bovine cells

We used commercial antibodies against PHB-1 protein (anti-PHB-1) for localization of host PHB-1. First, we analyzed the ability of the anti-PHB-1 to distinguish host PHB-1 and *Ta*PHB-1. We observed that the anti-PHB-1 could recognize host PHB-1 as a band at 30kDa was observed in western blot analysis with both BoMac cells and Ana 2014 cells (Fig 3A). Further, to confirm that anti-PHB-1 could recognize *Ta*PHB-1, we probed recombinant *Ta*PHB-1 with anti-PHB-1 antibodies. Anti-PHB-1 was able to recognize the parasite PHB-1 also (Fig 3B). Thus, we concluded that anti-PHB-1 recognizes both parasite and host PHB-1. We labelled host PHB-1 with PHB-1 antibodies for confocal microscopy while we used TOMM antibodies, a mitochondrial marker, to label the mitochondria. We observed that bovine prohibitin-1 was majorly localized with in the host cell nucleus in Ana2014 cells (Fig 3C). We could not observe any co-localization of PHB-1 with TOMM suggesting that bovine PHB-1 was not present in the mitochondria (Fig 3C) of Ana2014 cells. However, in the uninfected cells, i.e., BoMac cells, the bovine PHB-1 was majorly observed in the cytosol which co-localizes with TOMM (Fig 3C) suggesting that in the uninfected cells bovine PHB-1 is present in the mitochondria. Thus, we conclude that the *T. annulata* infection leads to the export of host PHB-1 to the host cell nucleus from the mitochondria. Further, we observed that the exported *Ta*PHB-1 was present in the host cytosol but did not localize with TOMM suggesting that the localization of the host and parasite PHB-1 is different in the *Theileria* uninfected and infected cells.

**Fig 3.**
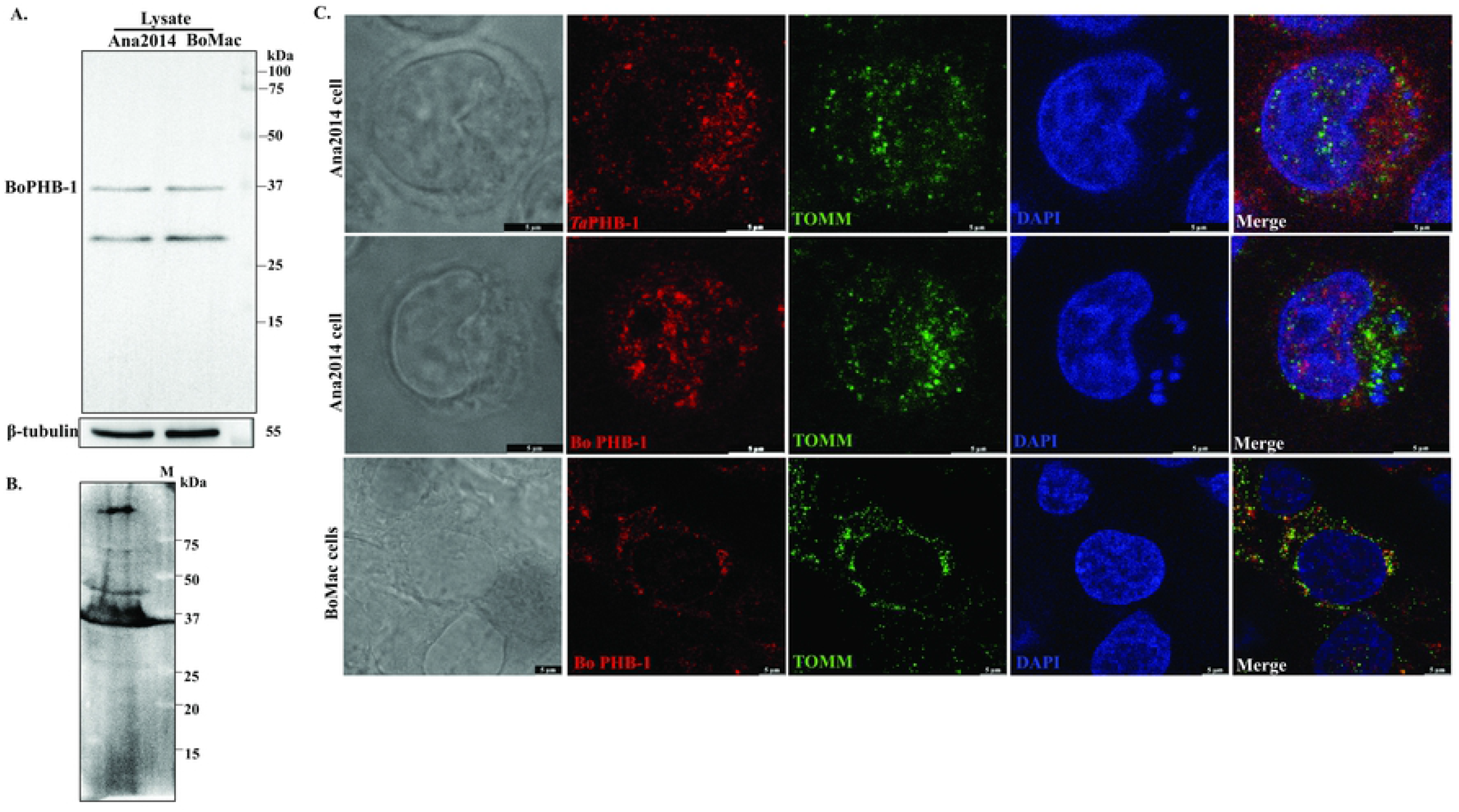
Sub cellular localization of *Ta*PHB-1 differs from bovine PHB-1 in Ana2014 cells. **A.** Ana2014 and BoMac cell lysates were probed with prohibitin antibody and β–tubulin was used as loading control. **B.** Western blotting of the bacterial lysate of recombinant *Ta*PHB-1 probed with commercial prohibitin antibodies. **C.** *Ta*PHB-1 localized majorly in the host cell cytoplasm and bovine PHB in host cell nucleus. TOMM was used to locate mitochondria. Scale bar-5 μm.

### *Theileria annulata* irreversibly changes the localization of BoPHB-1 in the infected cells

Ana2014 cells were treated for 72 h with Buparvaquone, a theileriacidal drug, to understand the role of parasite in changing in the localization of BoPHB-1. We observed inhibition of 70 % cell proliferation after the treatment (Data not shown). The quantification of transcripts for *Ta*SP, *Ta*PHB-1, and BoPHB-1 suggest decrease in the transcription with respect to bovine actin (Fig 4A). The effect of parasite on the localization of *Ta*SP, and BoPHB-1was further visualized using confocal microscopy. We observed until 72 h post treatment with Buparvaquone that there was no major change in the localization of the BoPHB-1 in the host cells while there was a significant decrease in the *Ta*SP (Fig 4B). This observation led us to hypothesize that the parasite makes irreversible changes in the localization of the BoPHB-1 in the transformed cells.

**Fig 4.**
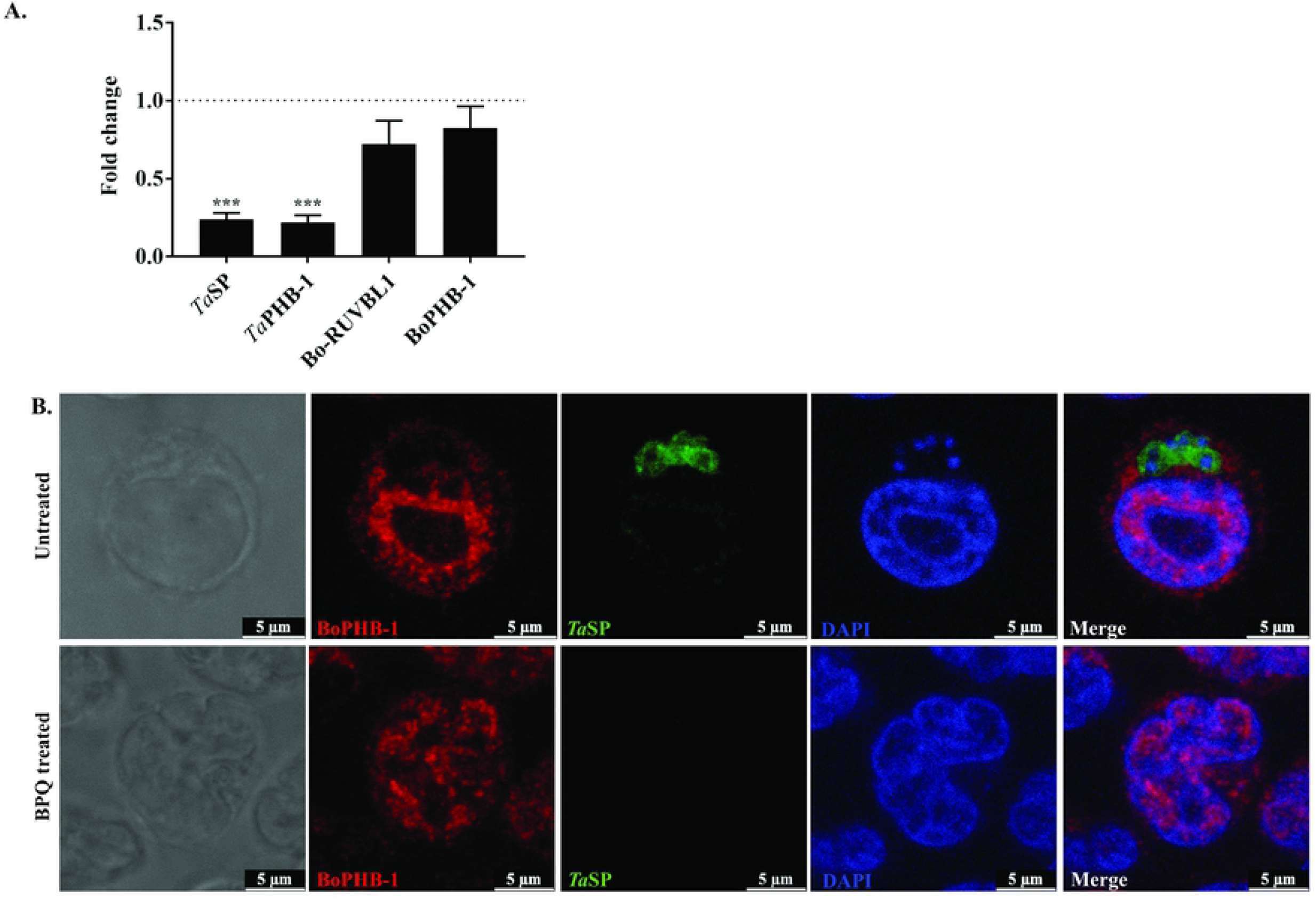
*Theileria annulata* irreversibly changes the localization of bovine prohibitin. **A.** Quantitative real-time PCR data showing the significant decrease in the level of transcripts of *Ta*SP and *Ta*PHB-1 upon buparvaquone treatment while there was no significant decrease in the transcripts of bovine RUVBL1 and bovine PHB-1. **B.** The localization of BoPHB-1 was unaffected in Ana2014 cells upon the elimination of parasite using buparvaquone. Untreated and BPQ treated cells were probed with BoPHB-1, *Ta*SP antibodies followed by staining with VECTASHIELD® antifade mounting medium with DAPI. Scale bar-5 μm.

### The yeast two-hybrid cDNA library (prey library) of Ana2014 cells contains sufficient number of independent clones

We prepared a yeast two hybrid library containing cDNA from the *T. annulata* infected bovine leucocytes (Ana2014 cells) to identify protein(s) that could interact with *Ta*PHB-1. One of the important criteria for an efficient cDNA library is that it should represent majority of the transcripts present in the cell. So, for preparing the cDNA library, we used cDNA from a range between 200 bp-2500 bp (S5 Fig). The titre obtained for the cDNA library with pGADT7-Rec in Y187 yeast strain was 2.4 × 10^6^ cfu that suggests that the library represents large number of independent clones. Thus, we used this library for yeast two-hybrid screening.

### The bait protein (*Ta*PHB-1) does not auto-activate reporter genes of Y2HGold yeast cells and is not toxic to the cells

Before mating of the prey and bait plasmids in yeast cell, we wanted to rule out the possibility of auto-activation and toxicity of bait plasmid in the Y2HGold yeast cells. Thus, the coding sequence of *Ta*PHB-1 was cloned in pGBKT7 plasmid. The western blot analysis using myc antibody detected a myc tagged *Ta*PHB-1 at the expected size of 50.3 kDa in Y2HGold yeast cells which confirmed the expression of this protein in the yeast cells (Fig 5A).

**Fig 5.**
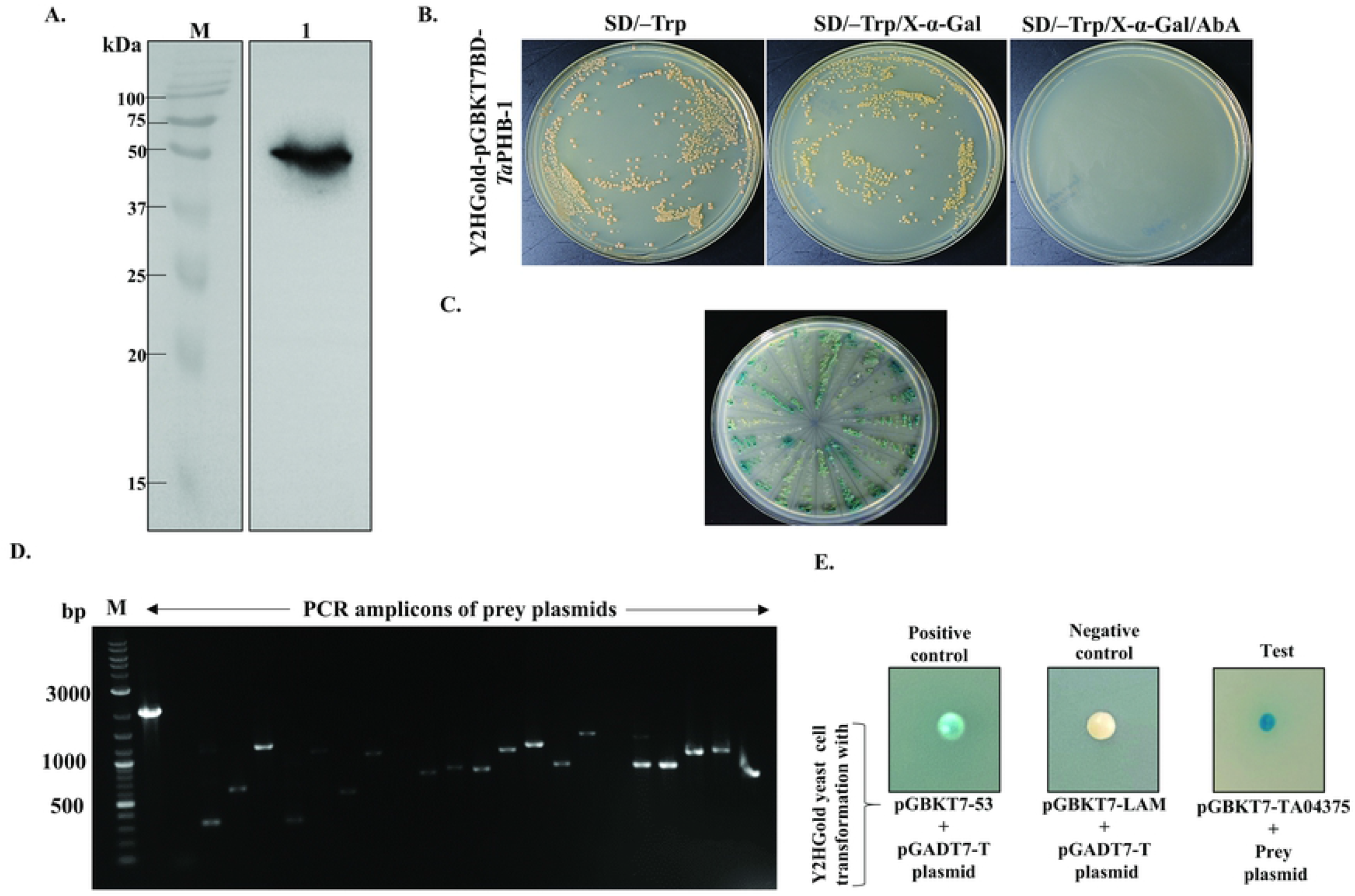
Yeast two hybrid library screening. **A** Western blot showing the expression of myc tagged *Ta*PHB-1 in Y2HGold yeast cells. **B.** White colonies on SD/-Trp/X-α-Gal agar plate and no colonies on SD/-Trp/X-α-Gal/AbA plates representing no auto-activation of reporter genes by TaPHB-1 in Y2HGold yeast cells. **C.** Plates showing Y2HGold cells that grew on QDO/X/A agar plates after library screening, **D.** Agarose gel electrophoresis analysis of PCR products of prey plasmids isolated from the colonies that grew on QDO/X/A agar plates. M: Marker, **E.** Growth of Y2HGold yeast cells on QDO/X/A agar plates after co-transformation of potential prey plasmid and pGBKT7-*Ta*PHB-1 confirmed their interaction. Co-transformation with pGBKT7-53 and pGADT7-T plasmids was used as a positive control and co-transformation with pGBKT7-LAM and pGADT7-T plasmids was used as a negative control on DDO/X/agar plate.

The pGBKT7-*Ta*PHB-1 construct was transformed into Y2HGold yeast strain to test the auto-activation of reporter genes expression by *Ta*PHB-1. The transformation mixture was spread on SD/–Trp, SD/–Trp/X-α-Gal and SD/–Trp/X-α-Gal/AbA agar plates. The presence of white colonies on SD/–Trp/X-α-Gal agar plate and the absence of yeast colonies on SD/–Trp/X-α-Gal/AbA agar plate after 3 days of incubation confirmed that there was no auto-activation of reporter genes by *Ta*PHB-1 in Y2HGold yeast cells (Fig 5B). Also, the size of the colonies on SD/–Trp agar plate were observed to be normal which suggests that *Ta*PHB-1 has no toxic effect on Y2HGold yeast cells.

### Yeast two-hybrid library screening led to the identification of bovine RUVBL1 as an interacting partner of *Ta*PHB-1

The yeast two-hybrid library screening was performed as mentioned in the methods section. Sixty yeast colonies grew after mating and selection of Y2HGold yeast cells containing *Ta*PHB-1 and cDNA library in the Y187 yeast cells on DDO/X/A agar plates. All these 60 colonies were patched on highly stringent QDO/X/A agar plates. Only 28 blue colonies out of these 60 colonies patched were able to grow on the QDO/X/A agar plates (Fig 5C). The colony PCR performed from each colony led to the identification of various duplicate colonies (Fig 5D). The size distribution of the insert in the prey plasmid is listed out in the S6 Fig. The prey plasmids with unique molecular sizes (PCR product) was rescued with ampicillin resistance marker in *E. coli* DH5α cells. All rescued prey plasmids were re-transformed along with pGBKT7-*Ta*PHB-1 into Y2HGold yeast cells and selected on QDO/X/A agar plates to rule out false positive in pervious steps. Only one prey plasmid out of all the plasmids was able to grow on the QDO/X/A agar plate after co-transformation (Fig 5E). The sequence analysis of confirmed prey plasmid using NCBI-BLAST tool led to the identification of the interacting gene as *Bos taurus* RuvB like AAA ATPase 1 (RUVBL1).

### *Ta*PHB-1 interacts with bovine RUVBL1 *in vitro*

The interaction of *Ta*PHB-1 and bovine RUVBL1 (Bo-RUVBL1) was reconfirmed by the analysis of interaction of these two proteins by co-expression *in vitro*. We used pETDuet-1 vector to co-express both the proteins in *E. coli* Lemo21(DE3) competent cells. The *Ta*PHB-1 was cloned into MCS1 of pETDuet-1 and full-length *B. taurus* RUVBL1 was cloned into MCS2 of pETDuet-1. Both proteins were found to express in the *E. coli* Lemo21(DE3) cells (S7 Fig). After induction with the IPTG, His tagged *Ta*PHB-1 was purified. The analysis of His-tag pull down of His-tagged *Ta*PHB-1 led to the identification of S-tagged Bo-RUVBL1 as a co-eluate which confirmed the interaction between *Ta*PHB-1 and bovine RUVBL1 (Fig 6A).

**Fig 6.**
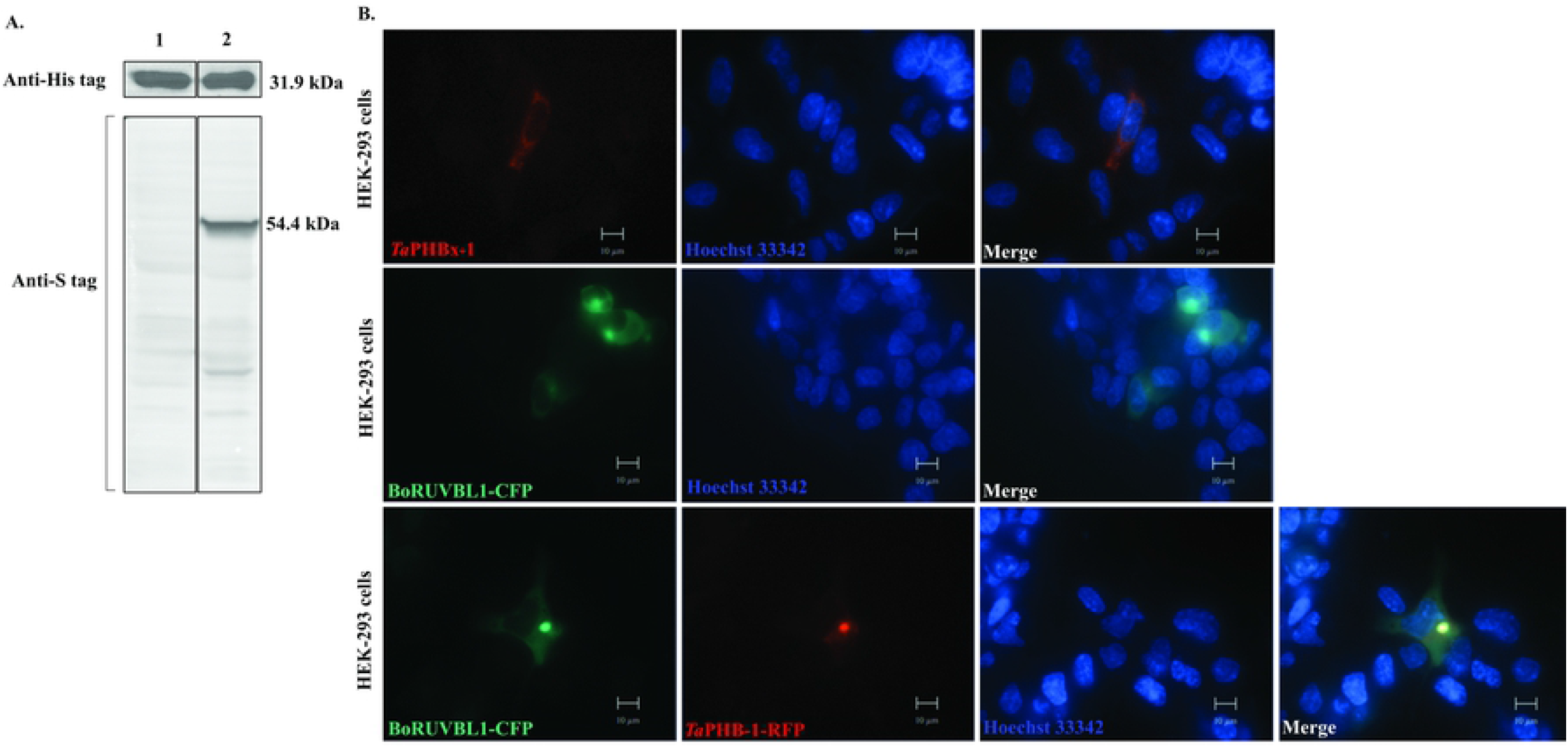
*Ta*PHB-1 interacts with bovine RUVBL1 *in vitro*. **A.** Western blot analysis of *Ta*PHB-1 (6xHis tag), RUVBL1 (S tag) after pulldown using Ni-NTA beads. 1: Eluate after pulldown of LemoDE3-pETDuet1-*Ta*PHB-1 bacterial cell lysate. 2: Eluate after pulldown of LemoDE3-pETDuet1-*Ta*PHB-1-RUVBL1 bacterial cell lysate. **B.** Colocalization of *Ta*PHB-1 (RFP tag) and bovine RUVBL1 (CFP tag) in HEK293 cells. HEK293 cells were transfected individually with pcDNA3RFP-*Ta*PHB-1 and pcDNA3CFP-BoRUVBL1 or co-transfected with both the plasmids using Lipofectamine® 3000 reagent. The fluorescence microscopy images were captured after 24 h after staining the nuclei with Hoechst 33342. Scale bar-10 μm.

The interaction between *Ta*PHB-1 and bovine RUVBL1 was also examined in HEK293 cells. Co-localization of RFP tagged *Ta*PHB-1 and CFP tagged bovine RUVBL1 in HEK293 cells further confirmed that *Ta*PHB-1 interacts with Bo-RUVBL1 (Fig 6B).

### *Ta*PHB-1 interacts with *B. taurus* RUVBL1 in Ana2014 cells

The interaction of *Ta*PHB-1 with Bo-RUVBL1 in Ana2014 cells was confirmed using the co-localization studies using antibodies specific to *Ta*PHB-1 and host RUVBL1. The bovine RUVBL1 was majorly localized in the host cell nucleus (Fig 7A and S8 Fig). The co-localization of *Ta*PHB-1 and Bo-RUVBL1 was observed majorly on the parasite surface (Fig 7A) which was localized using anti-*Ta*SP antibodies.

**Fig 7.**
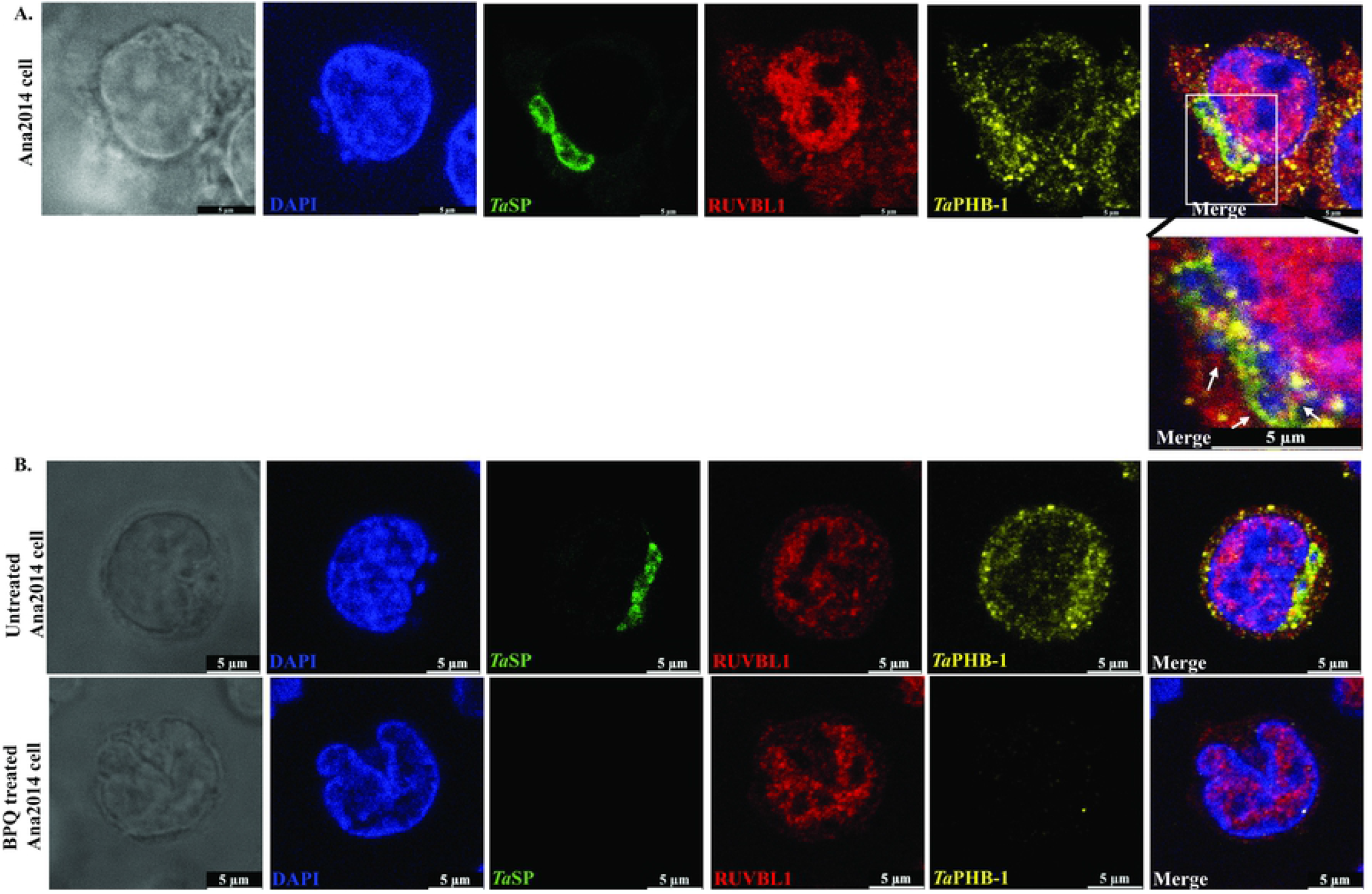
*Ta*PHB-1 co-localizes with bovine RUVBL1 on the surface of the parasite in Ana2014 cells. **A.** Ana2014 cells were immuno-stained with *Ta*PHB-1, *Ta*SP and RUVBL1 antibodies followed by staining with VECTASHIELD® antifade mounting medium with DAPI. Arrow indicates the site of co-localization of *Ta*PHB-1 and Bo-RUVBL1 on the parasite surface. **B. No effect on the localization of bovine RUVBL1 by elimination of the parasite.** The localization of Bo-RUVBL1 remain unchanged in Ana2014 cells upon the elimination of parasite using buparvaquone. Untreated and BPQ treated cells were probed with *Ta*SP, Bo-RUVBL1, *Ta*PHB-1 antibodies followed by staining with VECTASHIELD® antifade mounting medium with DAPI. Scale bar-5 μm.

Further, to analyze the effect of parasite on the localization of the BoRUVBL1, we treated Ana2014 cells with buparvaquone. The quantification of transcript for Bo-RUVBL1 showed that there was no significant decrease in the transcript of BoRUVBL-1 in buparvaquone treated Ana2014 cells (Fig 4A). Similarly, there was no significant change in the localization of Bo-RUVBL-1 upon killing of the parasite while there was decrease in the expression of parasite proteins, namely *Ta*SP and *Ta*-PHB-1 protein (Fig 7B).

As there are no reports of interaction between *Ta*PHB-1 and bovine RUVBL1 we speculated that the interaction between these two proteins might involve other interactions also. In order to identify the interactome, we performed co-immunoprecipitation using affinity purified *Ta*PHB-1 antibodies against the Ana2014 cell lysate. The bovine RUVBL1 was observed in the eluate through western blotting which confirms the interaction between *Ta*PHB-1 and bovine RUVBL1 (Fig 8A). Pre-immune sera was used as negative control in co-immunoprecipitation. Further, the eluate of co-immunoprecipitation was analyzed by mass spectrometry. The mass spectrometry led to the identification of 45 bovine proteins and 2 parasite proteins with more than ten unique peptides with less than 0.05 false discovery rate (Fig 8B, S2 sheet). We confirmed the result of LC-MS/MS by western blot analysis of one of the proteins, RPL7A, identified in the mass spectrometry analysis (Fig 8C). The interactome analysis of *Ta*PHB-1 by STRING suggests that the proteins involved in protein folding, cellular metabolic process, mRNA processing, catalytic activity, transporter activity and actin cytoskeleton organization are involved in the *Ta*PHB-1 interactome (Fig 8B).

**Fig 8.**
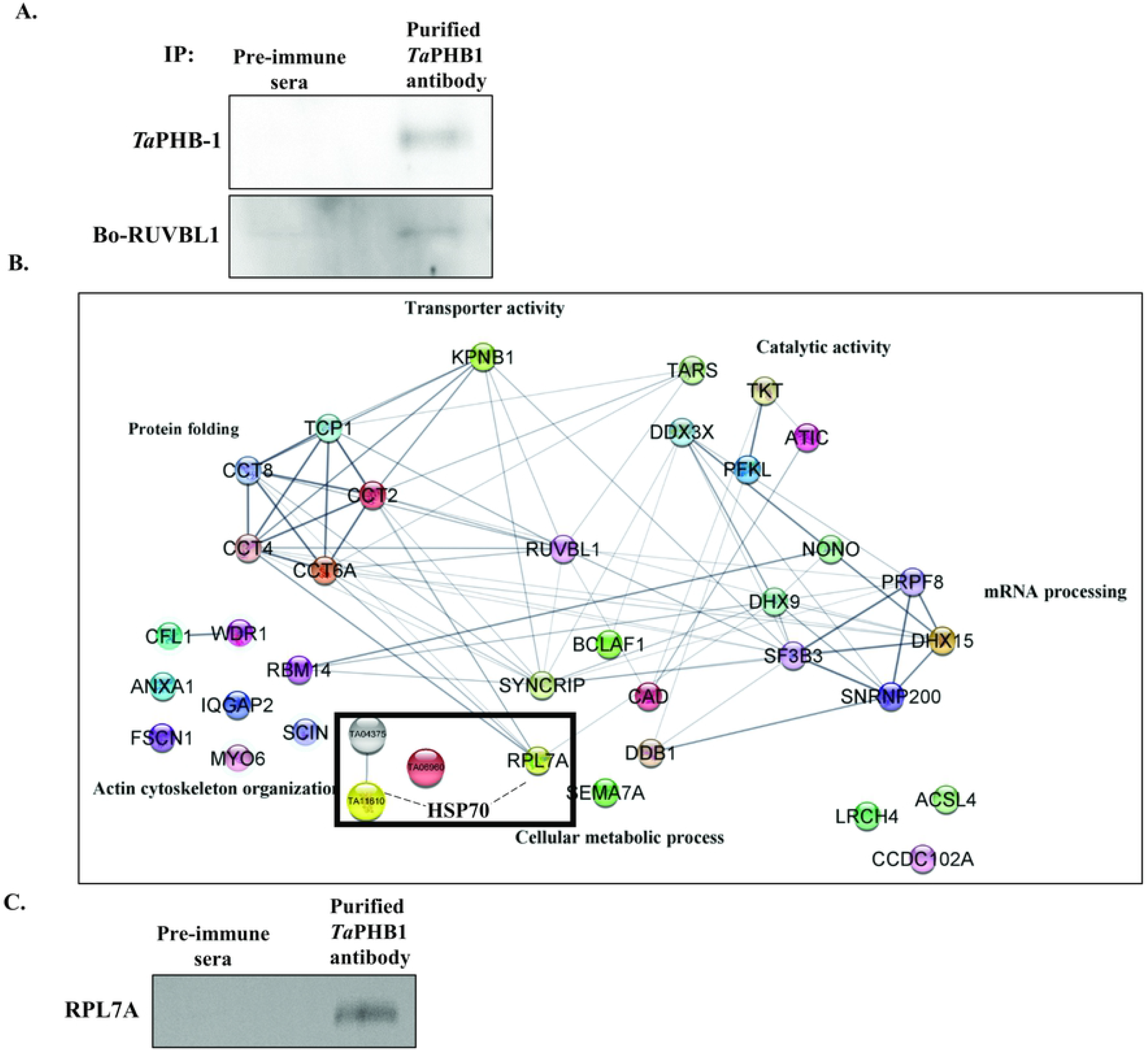
Co-Immunoprecipitation of Ana2014 cell lysate using *Ta*PHB-1 affinity purified antibodies and LC-MS/MS analysis of the interactome. **A.** Western blot analysis of eluate of co-IP of Ana2014 cells using affinity purified *Ta*PHB-1 antibodies. Pre-immune sera was used as control. **B.** STRING analysis of the proteins obtained in the *Ta*PHB-1 co-IP experiment (after subtraction with proteins obtained from pre-immune sera Co-IP). **C.** Confirmation of LC-MS/MS data. Western blot analysis of eluate of co-IP of Ana2014 cells using affinity purified *Ta*PHB-1 antibodies. RPL7A protein identified in the LC-MS/MS analysis was present in the pull down.

## Discussion

Some species of the *Theileria* parasite, such as *T. annualta* and *T. parva*, have a unique ability to transform the host cells. The sporozoites of the *Theileria* parasite can invade host leucocytes where they can control host cell proliferation. The presence of the parasite is essential for the transformation of the host cells as killing of the parasite inside host leads to the arrest in the proliferation of the host cell. This observation suggests that there must be parasite factors which could control metabolic/signaling pathways of the host cell. Hence, we quest for the identification of the parasite protein which may be secreted to host cytosol and may interact with the host protein(s). Our *in silico* analysis for such proteins helped us to shortlist few potential proteins. Previously, other research groups have also attempted to shortlist such proteins (8, 19). Prohibitins are one of the shortlisted proteins family. These are conserved proteins belonging to the family stomatin-prohibitin-flotillin-HflC/K (SPFH). They perform various cellular functions being localized to different cell organelles like nucleus, mitochondria and plasma membrane. Prohibitins are involved in the regulation of cell cycle, maintenance of mitochondrial membrane stability, control apoptosis, mitophagy, cell proliferation and also acts as molecular chaperone (23). Previously, parasite prohibitins have been assigned with divergent roles (16, 17, 24). We observed that the parasite prohibitins are quite divergent in comparison to mammalian prohibitins and most of the apicomplexans have one extra prohibitin. Our analysis suggested that *Ta*PHB1 out of three *T. annulata* prohibitins could be secreted out to the host cytoplasm. Thus, we focused our analysis on *Ta*PHB-1. As predicted, *Ta*PHB-1was observed on the parasite surface and in the host cytoplasm.

In normal cells the mammalian prohibitin 1 localizes to mitochondria and plays an important role in maintaining the stability of mitochondria by acting as a chaperone to safeguard newly imported proteins from m-AAA proteases (25, 26). Mammalian prohibitin was reported as a tumor suppressor with its anti-proliferative property (27). However, the oncogenic microRNA-27a which is upregulated in many cancers targets the 3’-UTR of prohibitin 1 and downregulates prohibitin 1 causing the cancer cell proliferation (27). In cancer cells, prohibitin 1 is mainly located in the nucleus (28) and co-localizes with proto-oncogenes c-myc, c-fos, p53 and Rb (29). In bovine macrophage cell line, i.e., BoMac, the bovine PHB-1 was found to be localized mainly in the mitochondria as it co-localizes with the mitochondrial marker, TOMM. However, in the case of Ana2014 cells, the localization host PHB-1 was observed mainly in the nucleus of the host cell. This observation again suggests that *Theileria* infected bovine leucocyte develops features common to the cancer cells (30). We further investigated whether this change could be reversed by specific elimination of the parasite in the Ana2014 cells. The specific killing of parasite in the host does not change the localization of BoPHB-1 back to the cytoplasm of the host cells which suggests that the parasite makes irreversible change in the localization of the Bo-PHB-1 in the host cells.

The observation that the *Ta*PHB-1 is secreted to the host cytoplasm led to speculate that it should be involved in protein-protein interaction with host protein(s). The yeast two-hybrid system is a well-established method for screening of interacting partners of interested protein (bait protein) against a library of proteins (prey library) (31). Thus we prepared a prey library with the cDNA of Ana2014 cells. The prey library which represents all the transcripts in the Ana2014 cells was used to screen the interacting partners of *Ta*PHB-1. The yeast two-hybrid screening studies led to the identification of bovine RUVBL1 as an interacting partner of *Ta*PHB-1. Further, to validate our results, we used other methods such as co-expression and pull-down methods. We used pETDuet1 plasmid to study the interaction between two proteins of interest *in vitro* (32). The co-expression of bovine RUVBL1 and *Ta*PHB-1 in bacterial system and pull-down of RUVBL1 with the *Ta*PHB-1 gave us further proof that these proteins interact with each other. Further, in order to validate the interaction of bovine RUVBL1 and *Ta*PHB-1, we co-expressed both the proteins in the HEK293 cells and performed co-localization studies. Both the proteins were found to co-localize. With these experiments we conclude that both these proteins interact with each other.

RUVBL1 (also known as pontin, TIP49) is an AAA+ ATPase and shares similarity with bacterial RuvB helicase (33). RUVBL1 is essential for cell proliferation (33, 34). RUVBL1 is over-expressed in various cancers (35–38) and downregulation or inhibition of RUVBL1 leads to controlled cancer cell proliferation (39, 40). RUVBL1 interacts with c-Myc and acts as transcriptional cofactor with ATPase and helicase activities (37). In nucleus, RUVBL1 interacts with telomere reverse transcriptase (TERT) which is essential for the maintenance of telomerase assembly (37). In the cytoplasm, RUVBL1 regulates Phosphatidylinositol 3-kinase-related protein kinase (PIKK) functions and involves in nonsense-mediated mRNA decay (41). No study till date has shown that RUVBL-1 can interact with PHB-1. The present study is the first ever report to identify the interactions between bovine RUVBL1 and *T. annulata* prohibitin (*Ta*PHB-1). Further, LC-MS/MS analysis deciphered the interactome of these two proteins. We speculate that *Ta*HSP70 is the link between bovine RUVBL1 and *T. annulata* prohibitin (*Ta*PHB-1).

In conclusion, we showed in the present study that the localization of bovine PHB-1 in *Theileria* infected bovine leucocytes changes from the host mitochondria to the host nucleus, while parasite prohibitin (*Ta*PHB-1) is exported out to the host cytoplasm and is also present on parasite surface. Further, we demonstrated that *Ta*PHB-1 interacts with bovine RUVBL1. The *Ta*PHB-1 interactome analysis suggests that *Ta*PHB-1 involves with those proteins which regulate actin cytoskeleton organization, protein folding, mRNA processing and metabolic process. We speculate that the interaction between PHB-1 and RUVBL1 will have implications not only in the arena of parasite biology but also in the field of cancer biology.

## Materials and Methods

### Ethical approval

The procedures for animal experiments were approved by the Institutional Animal Ethics Committee (IAEC) with approval no. IAEC/2021/NIAB/28/AS and performed in accordance with the guidelines from the Committee for the Purpose of Control and Supervision of Experiments on Animals (CPCSEA).

### Chemicals and Reagents

RPMI 1640 was purchased from Gibco, Life technologies. Fetal bovine serum was purchased from GE healthcare. Penicillin-streptomycin and TRIzol® Reagent were obtained from Invitrogen. Make Your Own “Mate & Plate™” Library System and Yeast Transformation System 2 were purchased from TaKaRa. Synthetic-defined (SD) broth/agar media, dropout (DO) supplements – Trp, -Leu, -Trp/-Leu, -Trp/-Leu/-Ade/-His, X-alpha-Gal (X-α-gal) and aureobasidin A (AbA) were obtained from TaKaRa. Plasmids, pcDNA3-CFP plasmid and pcDNA3-RFP were purchased from Addgene. Antibodies against His-tag and S-tag were purchased from Qiagen and GenScript respectively. Antibodies against RUVBL1 were purchased from Proteintech Group. Antibodies against β-tubulin, Prohibitin and myc-tag were obtained from Santa Cruz Biotechnology. Antibodies of TOMM (Abcam) were a kind gift from Prof. Naresh Babu V. Sepuri, University of Hyderabad, Hyderabad, India. Antibodies for RPL7A were purchased from Cell Signaling Technology. Secondary antibodies for microscopy, Alexa Fluor 647 conjugated anti-rabbit IgG, Alexa Fluor 555 conjugated anti-mouse IgG were purchased from Cell Signaling Technology. FITC conjugated anti-IgY was obtained from IgY immunologix Pvt. Ltd. Secondary antibodies for western blotting, anti-mouse IgG conjugated with HRP and anti-rabbit IgG conjugated with HRP were obtained from Pierce. VECTASHIELD antifade mounting medium with DAPI was purchased from VECTOR laboratories. Trypsin/LysC was purchased from Promega.

### Parasite and Cell culture

The in-house established *T. annulata* infected bovine leucocytes (Ana2014 cells) culture was maintained as described previously (42, 43). Briefly, *T. annulata* infected bovine leucocytes were cultured in RPMI 1640 (Gibco, Life technologies) reconstituted with 10% heat inactivated fetal bovine serum (GE healthcare) and 100 μg/ml penicillin-streptomycin (Invitrogen). Cells were cultured at 37°C with 5% CO_2_ atmosphere under humidified condition. Ana2014 cells were treated with buparvaquone at a concentration of 500 ng/ml for 72 h with the replacement of drug after 48 h. The BoMac cells, a kind gift from Dr. Judy Stabel, was also cultured in a similar way as Ana2014 cells. HEK293 cells were cultured in DMEM media reconstituted with 10% heat inactivated fetal bovine serum and 100 μg/ml penicillin-streptomycin at 37°C in a humidified 5% CO_2_ incubator.

### *In-silico* analysis of TA04375 (*Ta*PHB-1)

The SignalP 3.0 server was used for searching proteins with signal sequence in the proteome of *T. annulata* (44). The proteins expressed in the macroschizont stage were further filtered (20, 21). Prohibitin protein sequences of *T. annulata, Theileria parva, Plasmodium falciparum, Toxoplasma gondii*, *Cryptosporidium parvum*, *Babesia bovis*, *Bos taurus* and *Homo sapiens* were downloaded from UniProt KnowledgeBase (UniProtKB). The phylogenetic tree was constructed using the maximum likelihood method with bootstrap value of 100 in Mega 5 software (45).

The multiple sequence alignment of prohibitin sequences of *T. annulata* was performed using Jalview (46). The CELLO server was used for predicting the subcellular localization of these prohibitins (47, 48).

### Expression, purification and generation of polyclonal sera against *Ta*SP

The coding sequence of TA17315 (*Ta*SP) from 76 bp to 495 bp was amplified by PCR from cDNA of Ana2014 cells using specific primers, TASP-F and TASP-R (S1 table). The *Ta*SP was digested with NdeI and XhoI, and cloned into the corresponding sites of pET-28a(+) (Novagen). The recombinant construct was transformed into *E. coli* Rosetta (DE3) competent cells. The culture was induced with 0.5 mM isopropyl 1-thio--D-galactopyranoside (IPTG) and further grown at 37°C for 4 h. The recombinant His-tagged *Ta*SP was purified in native condition. Briefly, cells were pelleted down and lysed by sonication in lysis buffer (50 mM NaH_2_PO_4_ (pH-8), 300 mM NaCl, 10 mM imidazole) and the supernatant was collected by centrifuging at 8000 *g* for 25 min at 4°C. The supernatant was passed through the equilibrated Ni-NTA-agarose affinity resin (Qiagen). The resin was washed with wash buffer (lysis buffer with 40 mM imidazole) and the protein was eluted in the elution buffer (lysis buffer with 200 mM imidazole). The eluted protein was dialyzed overnight with the dialysis buffer 1 (50 mM sodium phosphate buffer (pH-5.8) with 40 mM NaCl) at 4°C. The dialyzed protein was further purified by ion exchange chromatography. The dialyzed protein was passed through the equilibrated Q-Sepharose Fast Flow beads (GE Healthcare). The beads were washed with dialysis buffer 1 with gradual increase in NaCl concentrations 40 mM, 60 mM, 80 mM and 100 mM NaCl. The bound protein is eluted with elution buffer (dialysis buffer 1 with 500 mM NaCl). The eluted protein was dialyzed overnight with the dialysis buffer 2 (PBS, pH-7.2) at 4°C. The purified protein was concentrated using 10 kDa centrifugal filter (Amicon®) to a concentration of 1 mg/ml. The recombinant *Ta*SP was used for raising polyclonal antibodies in chicken at IgY immunologix Pvt. Ltd.

### Quantitative real-time PCR

Total RNA was extracted from Ana2014 cells using NucleoSpin® RNA plus (Macherey-Nagel, Germany) according to the manufacturer instructions. The quality of RNA was assessed by using NanoDropTM 1000 spectrophotometer (Thermo Fisher Scientific Inc., USA). RNA with A_260/280_ (absorbance ratio) of 2.0 was used for cDNA preparation. The cDNA was prepared from 1 μg of total RNA using PrimeScript 1^st^ strand cDNA Synthesis Kit (TaKaRa, Japan) according to the manufacturer’s instructions. Quantitative real-time RCR (qRT-PCR) was carried-out using SYBR Premix Ex Taq™ (Tli RNaseH Plus) (TaKaRa, Japan) and CFX96TM real-time PCR system (Biorad, USA). Bovine actin was used as endogenous control. The list of used primers were listed in S1 table. The expression levels were calculated by 2^-ΔΔCt^ method, normalizing with the Ct values of the bovine actin.

### Construction of yeast two-hybrid cDNA library (prey library) of *T. annulata* infected bovine leucocytes

The total RNA was extracted from Ana2014 cells using TRIzol® reagent (Invitrogen) following the manufacturer’s protocol. One microgram of total RNA after DNase I (Invitrogen) treatment was used for cDNA library preparation using Make Your Own “Mate & Plate™” Library System (TaKaRa). All the procedures were followed according to the manual provided. Briefly, the first-strand cDNA was synthesized using SMART MMLV reverse transcriptase using CDS III primer also SMART oligo (S1 table), which finally results in known sequences at both ends of cDNA. Next, the cDNA was amplified by long distance PCR amplification using Advantage polymerase mix (TaKaRa). The amplified ds cDNA was purified using CHROMA SPIN TE-400 column which retains and traps the DNA molecules of less than 200 bp size. The ds cDNA after CHROMA SPIN TE-400 column purification was used for library construction by *in vivo* homologous recombination in yeast. The purified ds cDNA and SmaI linearized pGADT7-Rec were co-transformed into Y187 yeast strain using Yeastmaker Yeast Transformation System 2 (TaKaRa). The transformation mix was spread on to the SD/-Leu agar plates and incubated at 30°C for 3–4 days. All the transformants were harvested and pooled in freezing medium (YPDA/25% glycerol), aliquoted and stored at −80°C. Also, the number of independent clones was determined.

### Construction of bait plasmid

The total RNA was extracted from Ana2014 cells as mentioned above. The cDNA was synthesized using SuperScript™ First-Strand Synthesis System (Invitrogen) following the manufacturer’s protocol. *Theileria annulata* putative prohibitin (TA04375, GenBank accession no. XM_949999.1, *Ta*PHB-1) was amplified by PCR using its specific primers: TA04375-F and TA04375-R (S1 table). The PCR product was purified using PCR purification kit (Macherey-Nagel) and digested with NdeI and BamHI restriction enzymes. The digested *Ta*PHB-1 gene was ligated with pGBKT7 (NdeI and BamHI digested) vector plasmid. The recombinant pGBKT7-*Ta*PHB-1 was confirmed by sequencing.

### Expression of bait protein in Y2HGold yeast cells

The recombinant pGBKT7-*Ta*PHB-1 was transformed into Y2HGold yeast strain following the Yeastmaker Yeast Transformation System 2 (TaKaRa). The transformants were screened on SD/-Trp agar plate. A colony from SD/-Trp was inoculated in 5 ml of SD/-Trp broth and cultured for overnight at 30°C, 250 rpm. The overnight culture was added to 50 ml of YPDA and cultured at 30°C till the OD_600_ reached 0.4-0.6. The protein extract was prepared by lysing the yeast cells in cracking buffer as described previously (49). The protein lysate was resolved on 12% SDS-polyacrylamide gel and electro-transferred onto polyvinylidene fluoride (PVDF) membrane. After blocking the membrane in the blocking solution (3% skimmed milk powder in TBST), the membrane was incubated with myc antibody (Santa Cruz Biotechnology). The expression of myc tagged *Ta*PHB-1 was detected using horseradish-peroxidase (HRP) conjugated anti-mouse secondary antibody (Pierce) by SuperSignal West Pico chemiluminescent substrate (Thermo Scientific) in G: BOX Chemi imaging system (Syngene).

### Examining the autoactivation and toxicity of the bait protein in the yeast cells

The Matchmaker® Gold Yeast Two-Hybrid System (TaKaRa) protocols were followed to verify the autoactivation of reporter genes by *Ta*PHB-1. Briefly, the recombinant pGBKT7-*Ta*PHB-1 plasmid was transformed into Y2HGold yeast strain as mentioned above. The transformation mix was spread on SD/–Trp, SD/–Trp/X-α-Gal and SD/–Trp/X-α-Gal/AbA agar plates and incubated at 30°C for 3-5 days.

### Yeast two-hybrid library screening

The yeast-two-hybrid library screening was performed according to the protocols of the Matchmaker® Gold Yeast Two-Hybrid System (TaKaRa). Briefly, a single colony of Y2HGold-pGBKT7-*Ta*PHB-1 was inoculated in 50 ml of SD/–Trp broth medium and cultured for overnight at 30°C, 250 rpm. After the OD_600_ to ~0.8 reached, the yeast cells were pelleted down at 1000 g for 5 min and resuspended in fresh SD/–Trp broth medium so that the cell density reaches to 1×10^8^ cells/ml. The concentrated culture of bait was mixed with 1 ml of prey library culture and incubated at 30°C, 40 rpm. After 24 h, the yeast cells were pelleted down at 1000 g for 10 min, resuspended in 10 ml of 0.5X YPDA/kanamycin broth medium and cells were spread on to 150 mm SD/–Leu/–Trp/X-α-Gal/AbA (DDO/X/A) agar plates. Also, spreading on SD/-Trp, SD/-Leu and SD/–Leu/–Trp (DDO) agar plates was done to determine the mating efficiency. The agar plates were incubated at 30°C for 3-5 days. All the blue colonies which grew on DDO/X/A agar plates were patched on SD/-Ade/-His/-Leu/-Trp/ X-α-Gal/AbA (QDO/X/A) agar plate and incubated at 30°C for 3-5 days. The plasmids were extracted from all the colonies which grew on QDO/X/A agar plate using Easy Yeast Plasmid Isolation Kit (TaKaRa). PCR was carried out using the extracted plasmids as template, using forward primer (5’-TAATACGACTCACTATAG-3’) and reverse primer (5’-CTGTGCATCGTGCACCATCT-3’) specific to pGADT7Rec vector plasmid to identify similar size inserts.

### Confirmation of positive interactions and sequencing of the interacting partner of TA04375

The positive interactions were confirmed by co-transformation of bait and prey plasmids into Y2HGold yeast strain. After removing the duplicate prey plasmids based on size by PCR, each prey plasmid was transformed into *E. coli* DH5α competent cells (Invitrogen) and spread on LB/Ampicillin agar plates. The plasmids were extracted from the transformants using NucleoSpin plasmid purification kit (Macherey-Nagel). Each of these prey plasmids and previously constructed bait plasmids were co-transformed into Y2HGold yeast strain using Yeastmaker Yeast Transformation System 2 (TaKaRa). The transformation mix was spread on DDO/X and QDO/X/A agar plates and incubated at 30°C for 3-5 days. The plasmids of genuine positive interactions were sequenced. The prey plasmid was sequenced using the respective primers which were used for PCR. The sequence obtained was analyzed using BLAST tool of NCBI.

### *Ta*PHB-1-Bovine RUVBL1 interaction study

The coding sequence of *Ta*PHB-1 was PCR amplified from Ana2014 cells cDNA using its specific primers (S1 table). The *Ta*PHB-1 amplicons were digested with BamHI and SalI, and cloned into the corresponding sites on MCS1 of pETDuet-1 (Novagen) vector plasmid. Also, the coding sequence of *B. taurus* RUVBL1 was PCR amplified using Ana2014 cells cDNA and its specific primers (S1 table). The amplified RUVBL1 was digested with NdeI and BglII and cloned into the corresponding sites on MCS2 of pETDuet-1-*Ta*PHB-1.

The recombinant constructs (pETDuet-1-*Ta*PHB-1 and pETDuet-1-*Ta*PHB-1-RUVBL1 were separately transformed into *E. coli* Lemo21(DE3) (NEB) chemical competent cells. The cultures were induced with 0.4 mM IPTG and 1 mM rhamnose and further grown at 37°C for 4 h for the expression of recombinant proteins. The cells were pelleted down and lysed by sonication in lysis buffer (50 mM NaH_2_PO_4_ (pH-8), 300 mM NaCl, 10 mM imidazole). The cell lysate was incubated with Ni-NTA-agarose (GE Healthcare) affinity resin for overnight at 4°C. The resin was washed with 20 mM imidazole containing lysis buffer and finally the His-tagged *Ta*PHB-1 was eluted with 500 mM imidazole containing lysis buffer. The western blot analysis of the samples were performed as mentioned in the previous sections. The membrane was incubated with antibodies for His-tag-HRP or S-tag. The S-tagged bovine RUVBL1 was probed with anti-Rabbit-HRP secondary antibody.

### Co-localization studies

The full length *Ta*PHB-1 was PCR amplified from Ana2014 cells cDNA using the specific primers (S1 sheet). The amplified *Ta*PHB-1 was cloned into pcDNA3-RFP plasmid (#13032, Addgene) using HindIII and NotI restriction sites. Also, the full length bovine RUVBL1 was PCR amplified from Ana2014 cells cDNA using the specific primers (S1 table). The amplified RUVBL1 was cloned into pcDNA3-CFP plasmid (#13030, Addgene) using HindIII and NotI restriction sites. Both the clones were confirmed by double digestion and sequencing as well. The recombinant constructs were transfected into HEK293 cells either individually or together using Lipofectamine® 3000 reagent (Invitrogen) following the instructions provided. After 24 h of transfection, the fluorescence images of the HEK293 cells were captured using Axio Observer 7 microscope Apotome 2 (Carl Zeiss).

### Cloning, expression, purification of recombinant *Ta*PHB-1 and raising polyclonal antibodies

The coding sequence of *Ta*PHB-1 was PCR amplified from Ana2014 cells cDNA using specific primers (S1 table). The full length TA04375 was digested with NdeI and NotI, and cloned into the corresponding sites of pET-28a(+) (Invitrogen). The recombinant construct was transformed into *E. coli* Lemo21(DE3) competent cells (NEB). The culture was induced with 0.4 mM IPTG and 1 mM rhamnose and further grown at 37° C for 4 h. The recombinant His-tagged *Ta*PHB-1 was purified in denatured condition. Briefly, cells were pelleted down and lysed by sonication in lysis buffer (10 mM Tris-HCl (pH-8), 10 mM EDTA, 100 mM NaCl, 10 mM DTT) and the inclusion bodies were pelleted down by centrifuging at 13800 *g* for 25 min at 4°C. The inclusion bodies were washed with wash buffer 1 (50 mM phosphate buffer (pH-7.2), 10 mM EDTA, 200 mM NaCl, 2 M urea, 1% triton X-100) and subsequently with wash buffer 2 (50 mM phosphate buffer (pH-7.2), 1 M NaCl). The inclusion bodies were solubilized in solubilization buffer (10 mM Tris-HCl (pH-8), 100 mM NaH_2_PO_4_, 100 mM NaCl, 8 M urea) and, incubated with equilibrated Ni-NTA-agarose affinity resin for 2 h at room temperature. The resin was washed with solubilization buffer (pH-6.3). Recombinant *Ta*PHB-1 was eluted in solubilization buffer (pH-4.3). The eluted pure recombinant *Ta*PHB-1 was used for raising polyclonal antibodies in mice as described previously (50).

### Affinity purification of *Ta*PHB-1 mouse polyclonal antibodies

The antibodies specific to *Ta*PHB-1 were purified from mice sera by immunoblotting as described previously (51). Briefly, 500 μg of the purified *Ta*PHB-1 protein was separated on SDS-PAGE gel and, transferred onto PVDF membrane. The membrane was blocked with blocking buffer (5% BSA in TBST) for 2 h at room temperature. The protein band was stained by Ponceau S staining, the protein band was cut and washed with TBST until the membrane was destained. The membrane strip was incubated with 500 μl of mice sera for overnight at 4°C. The strip was washed thrice with TBST. Finally, the bound antibodies were eluted in 2 M glycine (pH-2.5) and the eluate was immediately neutralized with 1M Tris-HCl buffer (pH-9.0).

### Co-immunoprecipitation

Co-immunoprecipitation (Co-IP) was carried out using Pierce™ Direct Magnetic IP/Co-IP kit (Thermo Scientific) according the instructions provided by the manufacturer. Briefly, 10 μg of affinity purified *Ta*PHB-1 antibodies were cross-linked to the beads using Disuccinimidyl suberate (DSS). Ana2014 cells (40 × 10^6^ cells) were lysed in 1 ml of lysis buffer and incubated with the antibody cross-linked beads for overnight at 4°C. The beads were collected on a magnetic stand and washed thrice with the wash buffer. The bound proteins were eluted in the elution buffer and the eluate was neutralized with the neutralization buffer. The eluate was analyzed for the presence of both *Ta*PHB-1 and bovine RUVBL1 proteins by western blot analysis.

### Mass spectrometry analysis of Co-IP eluted proteins

The Co-IP eluate was processed for LC-MS/MS analysis by in-solution trypsin digestion method. The eluate was subjected to reduction with 20 mM DTT at 56°C for 1 h. Next, alkylation was carried out with 20 mM iodoacetamide (IAA) for 1 h at the room temperature in dark. The proteins were digested with trypsin/LysC (1:30 w/w) overnight at 37°C. The digestion reaction was stopped by adding 0.1% trifluoroacetic acid. The digested peptides were purified with C-18 spin columns. The purified peptides were concentrated by a vacuum evaporator and finally re-suspended in 0.1% formic acid. The peptides were analyzed by Q Exactive HF-Orbitrap mass spectrometer (Thermofisher Scientific) coupled with Ultimate 3000 RSLCnano LC system (Thermofisher Scientific). The peptides were injected into a reverse phase C-18 column (PepMap RSLC C18, 2 μm, 100 Å, 75 μm × 50 cm, Thermofisher Scientific) and separated by a gradient flow of solvent B (0.1% formic acid in 80/20 acetonitrile/water) from 5% to 90% in 180 mins. A parent ion scan was performed with a scan range of 375 to 1600 m/z with a resolution of 60,000. The top 25 intense peaks were fragmented by higher energy collision induced dissociation (HCD) fragmentation in MS/MS with the resolution of 15,000. The obtained spectra was analyzed using Thermo Proteome Discoverer software version 2.5. The database search was carried out using SEQUEST HT with the following parameters: maximum missed cleavage sites of 2, fragment mass tolerance of 0.02 kDa and precursor mass tolerance of 10 ppm. Fixed modification, carbamidomethylation of Cys (Cysteine) and dynamic modification, oxidation of Met (Methionine) were added. The resulting peptides were validated using percolator at a 5% False Discovery Rate (FDR), which validates using PEP (Psterior Error Probability) and q value. The interactome analysis was performed using STRING tool and Cytoscape ver 3.8.2 software.

### Immunofluorescence

The Ana2014 cells were washed twice with PBS and allowed to attach on poly-lysine coated cover slips. The cells were fixed and permeabilized in 3.7 % paraformaldehyde/ 0.18 % Triton X-100 (in PBS) at 37°C for 30 min. The Ana2014 cells were washed once with PBS and coverslip was blocked using blocking buffer (1% BSA in PBS) at 37°C for 1 h. The cells were incubated with the antibodies for RUVBL1, BoPHB-1, TOMM, *Ta*SP and affinity purified antibodies *Ta*PHB-1 at 4°C overnight. The cells were washed thrice with PBS and incubated with Alexa Fluor 647 conjugated anti-rabbit IgG (Cell Signaling Technology), Alexa Fluor 555 conjugated anti-mouse IgG (Cell Signaling Technology) and FITC conjugated anti-IgY (IgY immunologix) at room temperature for 1 h. The cells were washed thrice with PBS and the coverslip was mounted on a glass slide with VECTASHIELD® antifade mounting medium with DAPI (VECTOR laboratories). The images were captured using confocal microscope (Leica SP8, Leica Microsystems) and processed using LAS X software.

## Acknowledgements

We are grateful to Dr. Judy Stabel (Johne’s Disease Research Project, USDA-ARS-NADC) for providing BoMac cell lines. pcDNA3-CFP (Addgene plasmid # 13030) and pcDNA3-RFP (Addgene plasmid # #13032) was a gift from Doug Golenbock. We are grateful to Mr. Shashikant Gawai and Dr. Jayant Hole for their help in the confocal microscopy and the animal experiments respectively. We are thankful to Ms. S.V. Dilna and Ms. Nilanjana Ganguli for their help in LC-MS/MS analysis.

## Supplementary Figure

**S1 Fig.** Schematic representation of analysis of proteome for identification of proteins with potential to be involved in the transformation of the host cell.

**S2 Fig. *In silico* analysis of *T. annulata* prohibitins. A.** Multiple sequence alignment of *Ta*PHBs (Ta04375, Ta19320 and Ta08975) showing SPHF domain with a red color box and the amino acid having 100% identity are shown in color. **B.** Identity matrix of *Ta*PHBs showing that Ta08975 is highly divergent from Ta04375 and Ta19320.

**S3 Fig**. **Protein expression, purification, chicken polyclonal antibodies of *Ta*SP.** His tagged *Ta*SP protein purification using Ni-NTA agarose beads followed by ion exchange chromatography. M: Marker. 1: Coomassie brilliant blue stained gel-eluate with 500 mM NaCl during ion exchange chromatography, 2: Western blotting of purified *Ta*SP probed with His-tag antibody, 3: Western blotting of Ana2014 cell lysate probed with anti-*Ta*SP IgY antibodies.

**S4 Fig. LC-MS/MS analysis of recombinant *Ta*PHB-1.** LC-MS/MS analysis of recombinant *Ta*PHB-1 showing 69% coverage and 26 unique peptides. Peptides identified by LC-MS/MS are highlighted.

**S5 Fig.** Size distribution of ds cDNA of Ana2014 cells on 1% agarose gel. M: Marker and 1: Ana2014 cells ds cDNA.

**S6 Fig.** Size distribution of PCR products of prey plasmids using pGADT7-Rec specific primers.

**S7 Fig. Confirmation of expression induction of *Ta*PHB1 and bovine RUVBL1 in LemoDE3 bacterial cells.** Western blotting showing the expression of His tagged *Ta*PHB1 and S-tagged bovine RUVBL1 in LemoDE3 cells. M: Marker, 1: LemoDE3-pETDuet1-*Ta*PHB-1 bacterial cell lysate probed with His-tag antibodies. 2: LemoDE3-pETDuet1-*Ta*PHB-1-RUVBL1 bacterial cell lysate probed with S-tag antibodies.

**S8 Fig. Bovine RUVBL1 is not transported to the parasite in Ana2014 cells.** Bovine RUVBL1 was not observed in the parasite but only in the host cell nucleus and cytoplasm.

**S1 sheet.** Analysis of proteome for identification of molecules with potential to have role in transformation of host cell.

**S2 sheet**. List of proteins identified after Co-Immunoprecipitation of Ana2014 cell lysate using *Ta*PHB-1 affinity purified antibodies and LC-MS/MS analysis

**S1 table.** List of primers used in this study.

## Notes

### Competing Interest Statement

The authors have declared no competing interest.

